# EEG biomarkers of α5-GABA positive allosteric modulators in rodents

**DOI:** 10.1101/2024.03.26.586837

**Authors:** Frank Mazza, Alexandre Guet-McCreight, Thomas D. Prevot, Taufik Valiante, Etienne Sibille, Etay Hay

## Abstract

Reduced cortical inhibition mediated by gamma-aminobutyric acid (GABA) is reported in depression, anxiety disorders, and aging. Novel positive allosteric modulator that specifically target α5-GABA_A_ receptor subunit (α5-PAM), ligand GL-II-73, shows anxiolytic, antidepressant, and pro-cognitive effects without the common side effects associated with non-specific modulation by benzodiazepines such as diazepam (DZP), thus suggesting novel therapeutic potential. However, it is unknown if α5-PAM has detectable signatures in clinically-relevant brain electroencephalography (EEG). We analyzed EEG in freely moving rats at baseline and following injections of α5-PAM and DZP. We showed that α5-PAM specifically decreased theta peak power whereas DZP shifted peak power from high to low theta, while increasing beta and gamma power. EEG decomposition showed that these effects were periodic and corresponded to changes in theta oscillation event duration. Our study thus shows that α5-PAM has robust and distinct EEG biomarkers in rodents, indicating that EEG could enable non-invasive monitoring of α5-PAM treatment efficacy.

## Introduction

Cortical inhibition mediated by Gamma-aminobutyric acid (GABA) is reduced in depression^1,2^, anxiety disorder^3^, aging^4^, and neurodegenerative disorders^5^. Accordingly, nonspecific GABA_A_ receptor (GABA_A_-R) positive allosteric modulators in the benzodiazepine drug class, such as diazepam (DZP), have been used as common pharmacological treatments^6^. However, nonspecific potentiation by DZP leads to side effects, such as amnesia and sedation, limiting their therapeutic potential^7^. In contrast, studies in chronically-stressed rats, which exhibit many symptoms of depressions, showed that novel α-5 GABA_A-_R subunit positive allosteric modulators (α5-PAM) have anxiolytic, antidepressant, and pro-cognitive effects without common side effects associated with DZP, thus suggesting a novel therapeutic potential for depression and aging^8–15^. However, quantitative *in-vivo* measures of drug effects are needed for both development and clinical translation. Electroencephalography (EEG) offers a non-invasive and cost-effective method to monitor brain activity and treatment response, but it remains unknown whether α5-PAM effects exhibit distinct EEG biomarkers.

GABA_A_-Rs are a key component by which GABA exerts its inhibitory effect, whereby subunit composition determine channel localization and properties^16–18^. Whereas the α1 subunit is widely distributed across neuron and interneuron types and throughout the brain^19–21^, α5 subunits are primarily expressed in the apical dendrites of pyramidal neurons, and mainly in the hippocampus and neocortex^18^. α5-GABA_A_-Rs mediate lateral inhibition to pyramidal neurons by somatostatin-expressing (SST) interneurons targeting the apical dendrites^20,22–27^. Reduced α5-GABA_A_-Rs may underly deficits in GABAergic and SST signaling, which have been identified as contributors to depression and cognitive impairment^28,29^. α5 subunit knockout mice exhibit altered phasic and tonic currents, as well as cognitive changes including altered memory, executive function and fear conditioning^30–32^. Relatedly, silencing SST interneuron inhibition leads to depression symptoms^33^, which are reversed by α5-GABA_A_-Rs PAMs^33,34^.

α5-PAMs may have clear and distinct signatures in EEG due to their strong inhibitory modulation of the apical dendrites of L23 and L5 pyramidal neurons, which are the main contributors of the EEG dipole, thus offering a powerful tool for non-invasive treatment monitoring^35^. Previous works have extensively characterized the effects of benzodiazepines, which predominantly slow cortical theta (4 – 8 Hz) and increase beta (12 – 30 Hz) in a behavior-dependant manner, without affecting total cortical or hippocampal theta power^36^. Previous rat task studies show that EEG beta and gamma (30 – 50 Hz) power are uniquely elevated by GABA_A_-R α2/3-potentiating drugs, such as benzodiazepines^36^. It remains to be determined if α5-PAM displays distinct EEG signatures from DZP, such as involving power in theta band. Studies show that hippocampal SST interneuron stimulation slows theta oscillations^37^, and reduced SST inhibition decreases theta power in computational studies of detailed human cortical microcircuits^38^. Further, α5 subunit knockout mice show reduced hippocampal-dependant fear-associated learning specifically during theta stimulation, which are oscillations modulated by SST interneurons^39–42^.

In this study, we measured EEG in freely-moving rats that were administered α5-PAM ligand GL-II-73, which was previously shown to have antidepressant, pro-cognitive and anxiolytic effects in chronically-stressed mice^34^. We compared the EEG effect with both vehicle and DZP, and analyzed features of the EEG as putative biomarkers of α5-PAM compared to DZP. We then decomposed the PSD into aperiodic and periodic components to further differentiate EEG features following α5-PAM and DZP application. Finally, we compared the EEG features across different doses of each drug, relative to doses that were previously identified as optimal in preclinical mouse experiments^34^.

## Results

Male Sprague Dawley rats (*n* = 8) were injected with either vehicle, α5-PAM, or DZP in a random Latin-square design, with at least 1 week between injections (Figure 1A). EEG and EMG were recorded at baseline and following intravenous injection using surgically implanted single channel subcutaneous EEG and EMG sensors (Figure 1B-C). We analyzed active moving EEG by calculating the power spectral density (PSD). All baseline EEGs displayed a prominent theta peak (at 7.1 ± 0.4 Hz) and 1/f slope characteristic of wakefulness^43^ (Figure 1D), with no presentation of characteristic of NREM and REM sleep such as elevated delta or solely high theta^44^.

**Figure 1.**
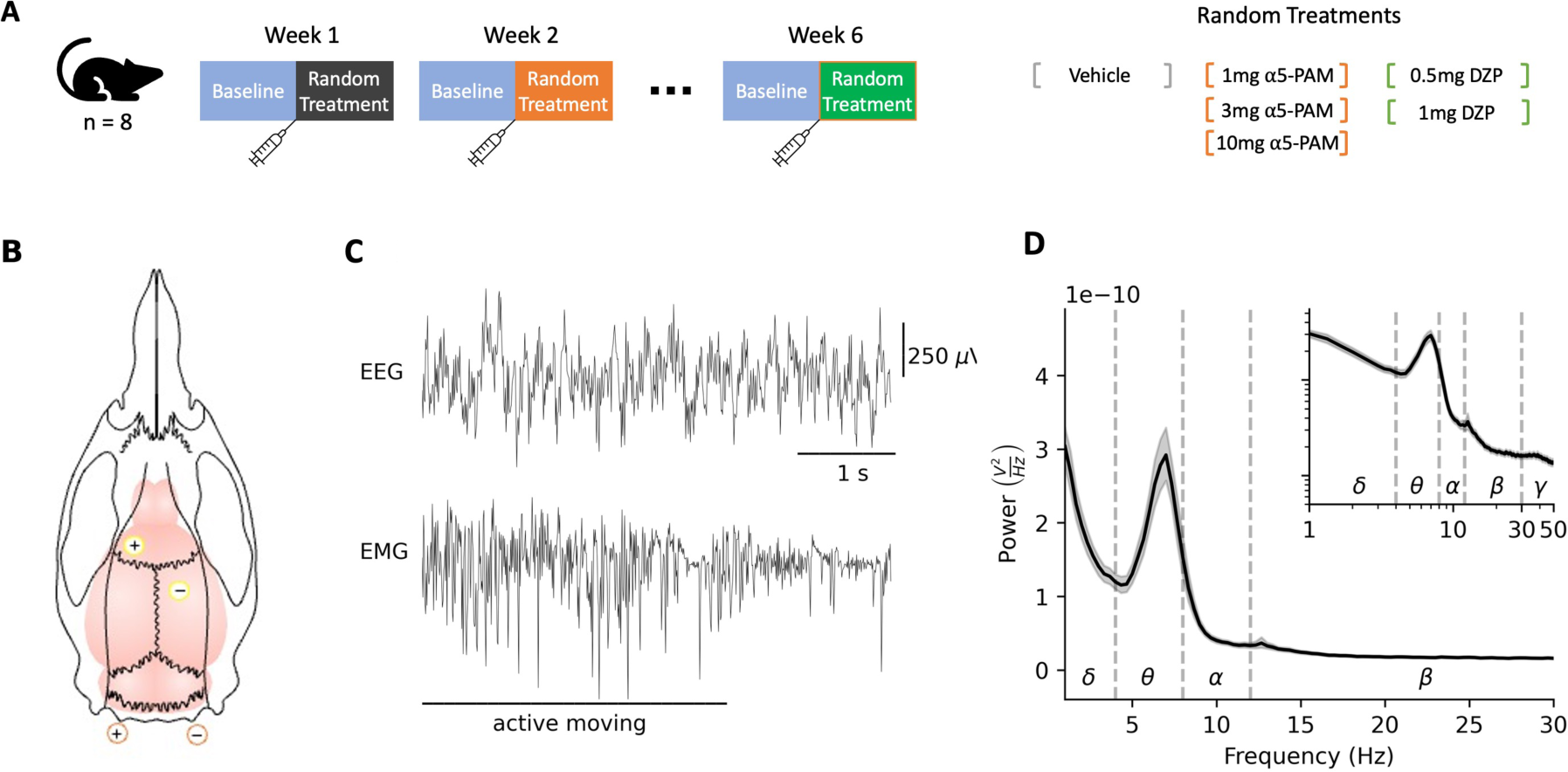
Testing a5-PAM and EEG recording in freely-moving rats. **(A)** Experimental protocol, where EEG and EMG were recorded for 60 minutes at baseline condition (pre-injection), and 60 minutes following injection of a random intravenous treatment, either vehicle, 1 mg α5-PAM, 3 mg α5-PAM, 10 mg α5-PAM, 0.5 mg DZP or 1 mg DZP. Rats were freely moving during the session. A week interval separated each injection session, so that at the end of 6 weeks, all types of injections had been administered. **(B)** Schematic showing subdural EEG electrode lead placement (yellow) and EMG electrode lead placement (orange). **(C)** Example of simultaneously recorded EEG and EMG while the rats were in active state. **(D)** Power spectral density plot of active-state EEG for baseline condition (n = 8 rats, bootstrapped mean and 95 % confidence intervals). Inset – same power spectral density plot shown on log-log scale.

We compared baseline and post-injection EEG for rats injected with vehicle, 3 mg α-5-PAM, and 1 mg DZP. Whereas EEG did not change significantly post-injection of vehicle (*p >* 0.05, Figure 2A), injection of α5-PAM decreased theta peak power in the raw averaged PSD (−36%, *p =* 0.046, *d* = −1.1, Figure 2B), and did not affect power in other frequency bands (all *p* > 0.05). In contrast, DZP injection tended to increase theta peak power but the statistical significance was at the trend level (+57%, *p* = 0.055), and DZP shifted the theta peak from high theta to low theta (increasing power in 4 – 5.3 Hz, and decreasing power in 6.6 – 8.6 Hz). DZP also increased beta power in 14 – 30 Hz (+160%, *p =* 0.001, *d* = 2.1) and increased gamma power in 30 – 44 Hz (+100%, *p <* 0.001, *d* = 2.4, Figure 2C). The window of 50 – 70 minutes post injection was chosen for analysis since it exhibited the largest and most consistent change in EEG spectrogram compared to baseline, in all injections (Figure 2G-I).

**Figure 2.**
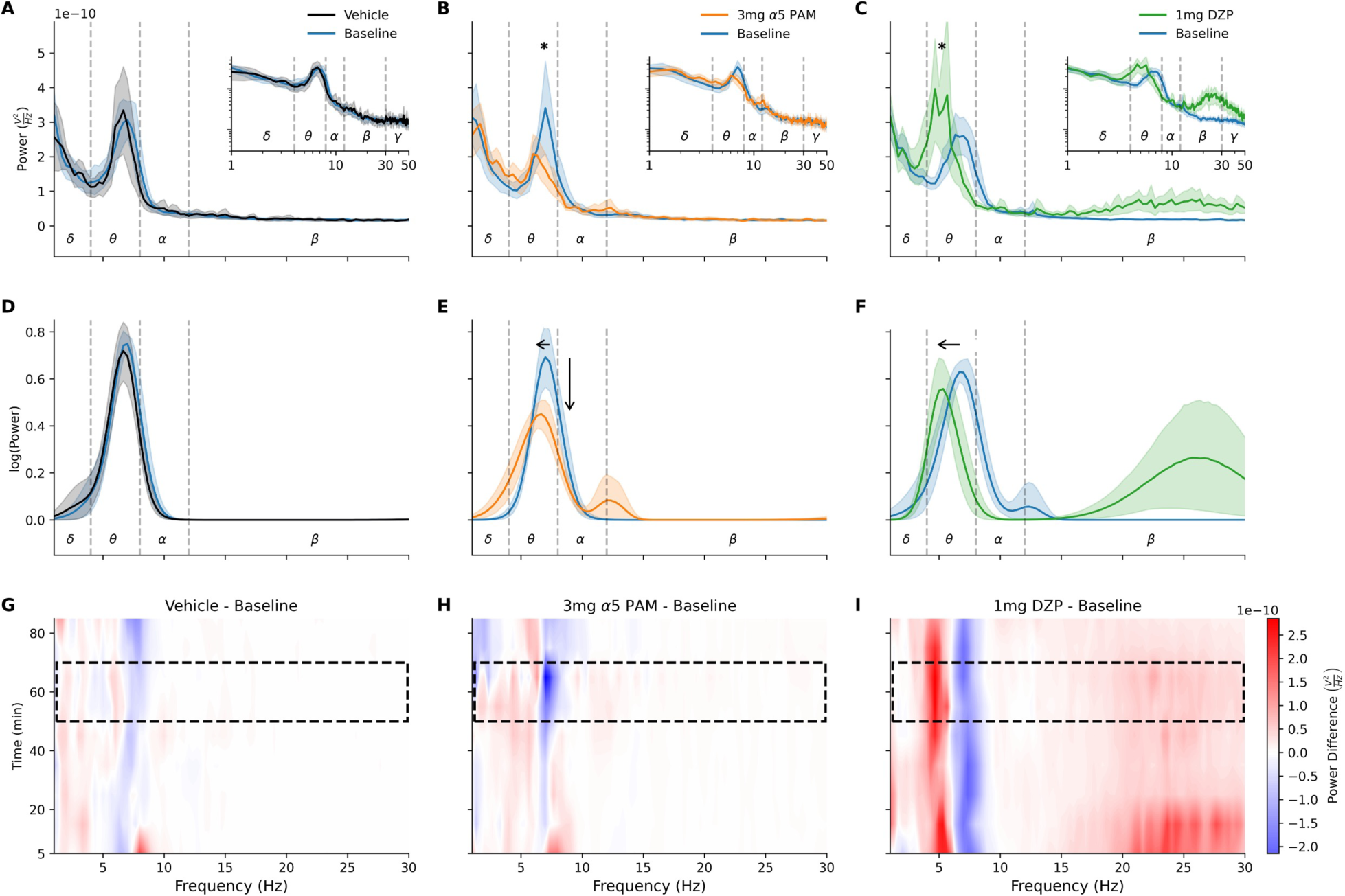
EEG signatures of α5-PAM in absolute and periodic component of PSD are specific from DZP. **(A)** PSD for active-state EEG recordings pre-injection (baseline, blue) and 50 – 70 minutes post-injection of vehicle (black, A), showing bootstrapped mean and 95% confidence intervals. Inset – same power spectral density data shown on log-log scale. **(B)** PSD in the case of 3 mg α5-PAM injection (orange) reduced peak theta power compared to baseline (blue). **(C)** PSD in the case of 1 mg DZP injection (green) did not reduce the peak theta power, but primarily left-shifted the peak and increased beta peak power. **(D)** Fitted periodic component for vehicle injection (black) and baseline (blue). **(E)** Fitted periodic component for 3 mg α5-PAM injection (orange) and baseline (blue) **(F)** Fitted periodic component for 1 mg DZP injection (green) compared to baseline (blue) **(G – I)** Spectrogram of difference from baseline in active-state EEG recordings 5 – 85 minutes post-injection of vehicle (G), 3 mg α5-PAM (H) or 1mg DZP (I). Dotted rectangle shows the window of 50 – 70 min used in post-injection PSD analysis for (A) – (F).

To better differentiate EEG changes due to α5-PAM versus DZP, we decomposed the PSD into aperiodic and periodic components and compared changes in periodic theta features (peak power, bandwidth, center frequency). α5-PAM injection decreased periodic theta power (−33%, *p* = 0.002, *d* = −1.6, Figure 2E), increased bandwidth (+35%, *p* = 0.005, *d* = 1.5), and slightly decreased the center frequency (−10%, *p* = 0.001, *d =* −2.0). In contrast, DZP did not significantly change periodic theta power (0%, *p* > 0.05, Figure 2F), but rather decreased bandwidth (−32%, *p* = 0.004, *d* = −1.5) and decreased center frequency (−18%, *p* = 0.001, *d =* −2.1) to a larger extent than α5-PAM (+14%, *p* = 0.001, *d =* −1.6). There were no significant changes in aperiodic component parameters, offset and exponent, for any conditions (all *p* > 0.05). Vehicle injection did not affect periodic theta power, bandwidth, or center frequency (all *p* > 0.05, Figure 2D).

To characterize the dose-response effect of α5-PAM and DZP on EEG, we analyzed additional doses of each drug (1 mg and 10 mg α5-PAM, as well as 0.5 mg DZP). The low and high doses of α5-PAM did not significantly affect the PSD, unlike the 3 mg α5-PAM dose (all *p* > 0.05, Figure 3 B, E, G). The lower dose of DZP showed a similar trend to 1 mg DZP, although to a smaller extent, by decreasing theta center frequency (−8%, *p* = 0.047, *d* = −0.9, Figure 3H), a borderline decrease of bandwidth (−20%, *p* = 0.120), and no change in theta peak power (*p >* 0. 05).

**Figure 3.**
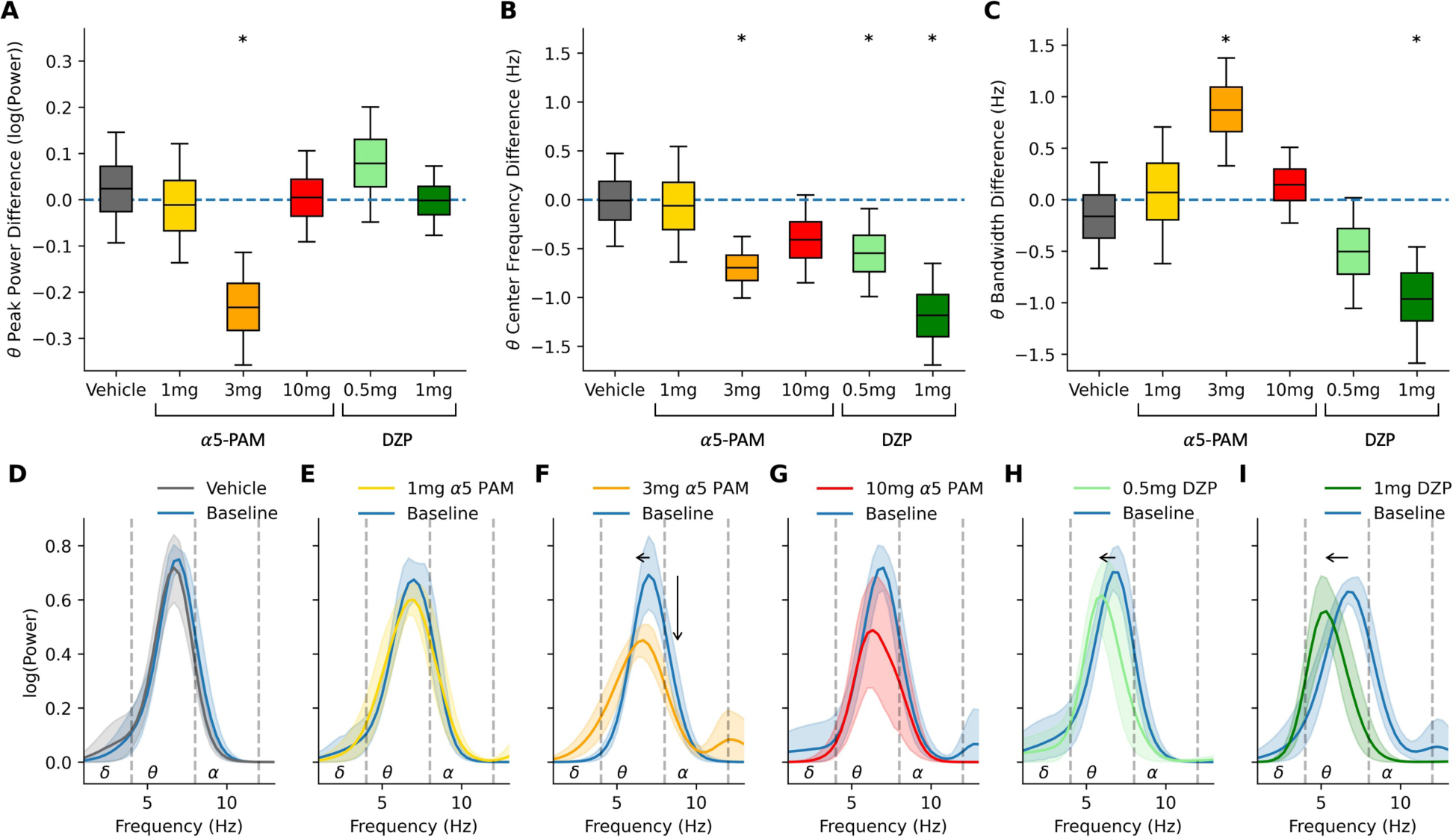
EEG signatures of different α5-PAM and DZP doses. **(A)** Periodic theta peak power (bootstrapped mean difference from baseline and 95% confidence interval) for the different conditions. Asterix shows significant difference between baseline and injection. **(B)** Same as (A) but for periodic theta center frequency. **(C)** Same as (A) but for periodic theta peak bandwidth. **(D– I)** Fitted periodic component of PSD pre-injection for baseline (blue) and 50 - 70 minutes post-injection of (D) vehicle, (E) 1 mg α5-PAM, (F) 3 mg α5-PAM, (G) 10 mg α5-PAM, (H) 0.5 mg DZP, (I) 1 mg DZP. Showing bootstrapped mean and 95 % confidence intervals. Vertical arrow shows significant difference in theta power peak, horizontal arrows show significant difference in theta power center frequency.

Finally, to determine the aspects of the EEG signal underlying the theta power modulation by the compounds, we detected low theta (4 – 6 Hz) and high theta (6 – 8 Hz) oscillatory events and compared their amplitude, frequency (events per second), and duration in rats at baseline and post-injection. α5-PAM decreased high-theta event duration compared to baseline (−31%, *p* < 0.001, *d* = −2.8) and slightly increased low-theta event duration (+9%, *p* = 0.004, *d =* 1.59, Figure 4C). DZP similarly decreased high theta event duration (−27%, *p* < 0.001, *d* = 2.3), but increased low theta event duration to a larger extent (+62%, *p* < 0.001, *d =* 2.1). DZP also decreased both low theta event frequency (−18%, *p* = 0.001, *d* = −2.2, Figure 4D) and high theta event frequency (−19%, *p* = 0.002, *d* = 1.7). Neither α5-PAM nor DZP affected event amplitude (Figure 4B), and vehicle did not affect any of the EEG waveform components.

**Figure 4.**
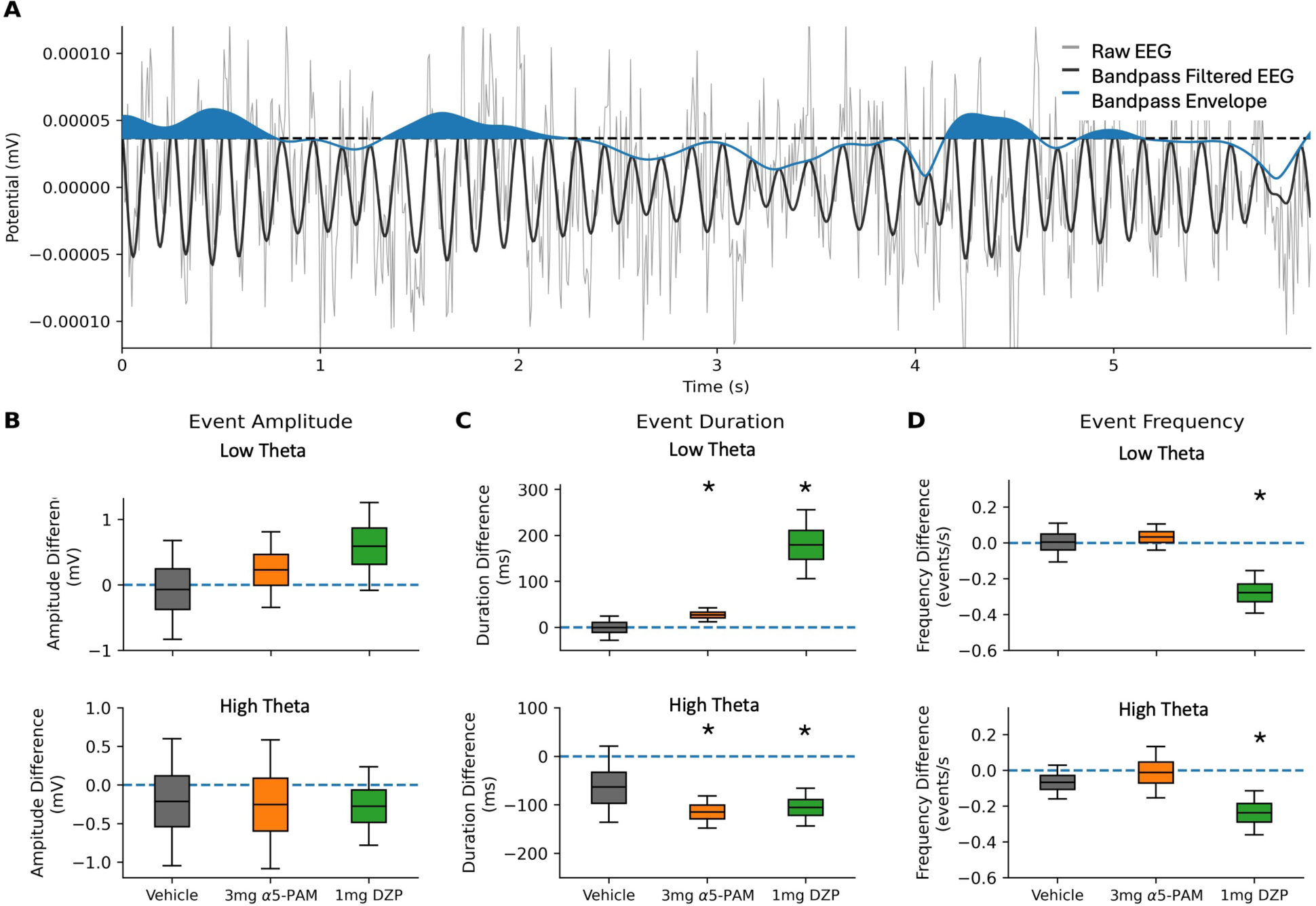
Signatures of α5-PAM and DZP in EEG waveform events. **(A)** Example trace of raw EEG waveform (grey), its high theta bandpass transformation (black), and instantaneous envelope (blue). Shaded blue area represents periods of theta events (elevated high-theta amplitude). Horizontal dotted line is the threshold for a large amplitude event. **(B)** Event amplitude in different conditions for low-theta (top) and high-theta (bottom) bandpass-filtered EEG (**C)** Same as (B) but for event duration. **(D)** Same as (B) but for event frequency. Error bars show bootstrapped mean difference from baseline and 95% confidence interval.

## Discussion

In this work, we identified in freely moving rats the EEG signatures of α5-PAM. Using GL-II-73, a ligand shown previously to have therapeutic effects in chronically-stressed mice in terms of mood and cognitive symptoms. We showed that α5-PAM had a distinct effect of decreasing peak theta power, whereas non-specific modulation by DZP shifted peak power from high theta to low theta. EEG decomposition and event analysis showed that these effects were seen more strongly in the periodic component of the decomposed PSD, and corresponded to decreased duration of EEG events in the theta band. We characterized the dose-response effect of α5-PAM on EEG and found that the dose of 3mg/kg had the largest specific effect on peak EEG power in the theta frequency band. Our study thus demonstrates that subunit-specific potentiation of α5-GABA_A-_R has distinct signatures in rodent EEG, suggesting candidate biomarkers for efficacy and monitoring of α5-PAM treatment for depression in human subjects.

The α5-PAM ligand GL-II-73, which was previously shown to have anxiolytic, antidepressant and pro-cognitive effects^34^, primarily decreased EEG theta power in this study. This is in line with increased resting-state theta power often seen in depression patients and that correlates with depression severity^45,46^. Increased theta power is also correlated with first-line treatment resistance and with second-line treatment response^45–50^, especially second-line treatments that target frontal cortical inhibition and decrease theta power following treatment^51,53–56^. Increased theta power has also been associated with cognitive impairement^57^, and increased theta coherence is associated with anxiety^58^. The decreased theta effect by the α5-PAM ligand reported in this study suggests that the inhibitory mechanisms it targets may underlie the EEG effects described above. This is supported by previous works indicating that reduced SST interneurons inhibition (which targets α5-GABA_A_-R^33^) in frontal regions is central to depression etiology^25,59,60^, and studies showing that SST interneurons provide rhythmic theta inhibition^61^. The link between reduced SST interneuron inhibition and theta power is further supported by detailed simulations of depression microcircuits showing that reduced SST interneuron inhibition primarily modulated theta power^62^, and by *in vivo* studies in the cortex and the hippocampus showing that theta oscillations are directly modulated by optogenetic stimulation of SST interneurons^37^. Altogether, these previous studies and our current results indicate that the theta modulation relevant to depression and treatment is likely mediated through α5-GABA_A-_R innervation by SST interneurons.

Although our study is the first to identify EEG biomarkers of α5 positive modulation, previous studies have identified EEG biomarkers of other GABA_A-_ Rs. For example, GABA α2/3 subunit potentiation has shown to increase beta and gamma power, consistent with benzodiazepine changes seen in previous studies^63^ and in line with diazepam effects we observed. As well, zolpidem, a potentiator of α1, was shown to modulate cortical and hippocampal delta power^36^. Together, our findings further support that specific GABA_A_-R subunit potentiation has signatures detectable in EEG. The robust EEG signatures of α5-GABA_A_-R potentiation are likely due to α5 subunit mediating inhibition onto layer 2/3 and 5/6 pyramidal neurons, which are the primary contributors to EEG, and more specifically due to the location of α5-GABA_A_-R on the apical dendrites, which strongly affect the EEG dipole^29,40^.

We identified global changes in theta power as a biomarker of α5-PAM, which constitute promising pre-clinical EEG correlates for treatment monitoring. Since a bipolar montage was used, where the positive electrode was placed atop the left frontal cortex and the negative reference electrode atop the right parietal cortex, our results reflect changes in relative activity and thus potential between these two regions^64^. The changes may thus reflect local increased frontal theta activity as seen in depression^51^, altered communication between frontal and hippocampal regions which is key for working memory^65^, or a combination of the two. These possibilities are in line with α5 subunits being abundantly expressed in rodent frontal and hippocampal regions, where they mediate SST interneuron inhibition^66^. Future studies will do well to include higher density EEG recordings to identify location-specific changes, and thus enhance the translation to human EEG. Relatedly, work to identify changes in theta power in other brain states will provide additional important biomarkers of α5 potentiation, as SST interneuron inhibition has been shown to play a role at rest^67^, during task^68^ and slow-wave sleep^69^. Previous *in vivo*^37,42^ and *in silico*^62^ work showing that SST interneurons modulate theta power during both task and at rest support the possibility of α5-PAM EEG signatures for additional states. Finally, future work should characterize the EEG signatures of α5-PAM in stressed rats to relate the antidepressant effects of α5-PAM to EEG biomarkers.

This is the first study to identify EEG signatures of α5-PAM ligand GL-II-73, which has been previously shown to have antidepressant, pro-cognitive, and anxiolytic effects in preclinical tests in chronically-stressed rodents^34^. The EEG signatures can serve as biomarkers for non-invasive monitoring of α5-PAM treatment efficacy in preclinical testing, to facilitate translation of the pharmacology to human trials. The similarity to EEG changes in depression indicate that the EEG biomarkers we characterized will be also relevant to monitoring the pharmacology efficacy in humans.

## Methods

### Animals

Sprague Dawley rats were provided by Charles River Laboratories (n = 10, age 7.3 ± 0.6 weeks, all male). The animal experiments, including EEG recordings, were performed at Charles River Laboratories. n = 8 rats were used at any given session of the study, since 2 rats were substituted with rats of similar age due to tail lesions found in weeks 4 and 6 of the study. Rats were group-housed in polycarbonate cages (2 – 3 per cage) and acclimated for at least 4 days in a 12-hour light/dark cycle with consistent room temperature (22 ± 2 C), 50% humidity, food and water *ad libitum*, as well as a nylon bone (Bio-Serv®, K3580) and tunnel retreat (Bio-Serv®, K3245) for enrichment. Surgery was performed to implant electrodes for EEG recording (see below). Rat age was 10.1 ± 0.2 weeks when EEG recordings started. All experiments were conducted in accordance with protocols approved by the Institutional Animal Care and Use Committee of Charles River Laboratories SSF.

### Surgery

EEG/EMG subcutaneous transmitters (model number: HD-S02) were obtained from Data Sciences International. Rats were anesthetized with isoflurane (2%, 800mL/min O2). Local anesthesia was provided with bupivacaine, and post-operative analgesia was provided with carprofen. The EEG transmitter was inserted into a pocket made close to the dorsal flank. Positive and negative leads were tunnelled subcutaneously towards the head. The positive lead was placed 2 mm anterior from the bregma and 2 mm lateral from the midline (left frontal cortex). The negative lead was placed 2 mm anterior to the lambda and 2 mm lateral from the midline (right parietal cortex). To record EMG activity, the positive lead and negative leads were sutured on the left and right musculus cervicoauricularis. Following surgery, rats were individually housed in cages. They were provided food and water *ad libitum*, as well as a nylon bone (Bio-Serv®, K3580) and paper towel for enrichment. Rats received a 5 - day course of antibiotics immediately following surgery. Pain was managed for 3 days. EEG recording began after a minimum of 10 days recovery following surgery.

### Injected Ligands

Rats were tested with one of 6 injected ligands from three groups - control vehicle, GABA receptor subunit α5 positive allosteric modulator (α5-PAM) of either 1, 3, or 10 mg dose, and diazepam (DZP) of either 0.5 or 1 mg dose (supplied by Sigma, Lot# 105F0451, purity > 99%). All ligands were held at room temperature. Ligands were formulated on days of use in a vehicle solution of 85% distilled H_2_O, 14% propylene glycol, and 1% Tween-80 for vehicle and α5-PAM, or 50% distilled H_2_O, 40% propylene glycol, and 10% alcohol for DZP. The route of administration for all ligands was through intravenous tail vein.

### Experimental design

Rats were injected with one of six test ligands in a standard Latin-square design. Rats were placed on RPC-1 receivers the night prior to recording for acclimation. Rats were recorded weekly (at least 5 days between each session). Baseline recording began at approximately 9am. After approximately 1 hour, rats were injected with a test ligand. Recording continued for an additional 24 hours following injection. All rats were euthanized via CO2 asphyxiation following experiment completion (Week 6). Video was recorded during the entirety of baseline and treatment.

### EEG Preprocessing

EEG and EMG data were down sampled to 140 Hz and visually inspected for artefacts. Periods of freely moving rats were identified by referencing temporally aligned video and electrophysiological recording, as well as the presence of EEG/EMG waveforms characteristic for this state. Periods were identified as freely-moving after at least 3 seconds of continuous moving data, excluding a 2 second buffer between freely-moving and other states (such as sleep). 60 minutes of baseline EEG and 50 – 70 min post-injection were included in the analysis. In 6 of the 48 sessions, where no high-quality moving EEG was found during this window, the window of inclusion was expanded to 30 – 70 min. The overall EEG length characterized as freely-moving was 15.2 (CI = 1.8 - 43.2) min for baseline and 4.8 (CI = 0.3 - 14.1) minutes for post-injection EEG. In 4 of 48 sessions the EEG length was less than 1 minute but it was sufficient to calculate the features of interest and it did not deviate from the group statistics.

### EEG Analyses

EEG power spectral density (PSD) were calculated using Welch’s method, with a 3 s Hanning window and 30% window overlap. We decomposed EEG PSDs into periodic and aperiodic components using FOOOF. The aperiodic component of the PSD was a 1/f function, defined by a vertical offset and exponent parameter. After removing the aperiodic component from the PSDs, we derived from the flattened spectrum the periodic components (representing putative oscillations) using gaussians, defined by center frequency (mean), bandwidth (variance), and power (height). Power spectra were fit across the frequency range 1 - 70 Hz with a resolution of 0.33 Hz. Fitting algorithm settings were as follows: width limits = [1.5, 8]; max number of peaks = 5; minimum peak height = .3; peak threshold = 3; and aperiodic mode = ‘fixed’, where the aperiodic component was fit as a straight line to the PSD. Canonical EEG bands were identified as delta (1 – 4 Hz), theta (4 – 8 Hz), alpha (8 – 12Hz), beta (12 – 30 Hz), and gamma (30 – 50 Hz). Group-wise differences across PSD curves were included in the analysis if a significant change in power was detected over a continuous span of 0.66 Hz.

### EEG theta event analysis

We analyzed characteristics of events of large amplitude in low theta (4 – 6 Hz) and high theta (6 – 8 Hz) to identify which aspect of EEG waveforms contributed to overall power changes in periodic theta. The threshold for a large-amplitude event was calculated by aggregating the average instantaneous envelope power (Hilbert transform) for baseline recordings. We calculated maximum envelope power and duration at sections for large-amplitude events. Event frequency was calculated as number of events per second. The same methodology was applied to 50 – 70 minute post-injection EEG, using the threshold established in the baseline EEG.

### Inclusion / exclusion criteria

3 of the 48 recording sessions were removed from analysis if the EEG exhibited outlier behaviour using the standard Q3 + 1.5x IQR method referencing peak theta power.

### Statistical Tests

To provide a more robust estimate of group differences despite the small sample size, a significant difference between two groups was determined if the 95% confidence intervals of their bootstrapped difference did not include 0 (10,000 bootstrap iterations). Between-group comparisons were achieved using the same method, and Bonferroni corrected for multiple comparisons.

## Acknowledgements

FM, AGM and EH thank the Krembil Foundation for their generous funding support. FM was also supported by Ontario Graduate Scholarship.

